## Supplementary Materials for "Large-scale gene losses underlie the genome evolution of parasitic plant *Cuscuta australis*" (from another version of this preprint: version 1, 2018-03-20)

**Description of Supplementary Files**

File Name: Supplementary Information

Description: Supplementary Figures, Supplementary Tables, Supplementary Notes and Supplementary References

File Name: Supplementary Data 1

Description: Supplementary Data 1a. Gene family expansion and contraction;

Supplementary Data 1b. Number of orthogroup genes in each species;

Supplementary Data 1c. Gene loss identified using Arabidopsis genes as reference;

Supplementary Data 1d. Tissue-specific expression levels of the orthogroup genes in *Solanum lycopersicum*, *Ipomoea nil*, *and Cuscuta pentagona*

File Name: Supplementary Data 2

Description: Supplementary Data 2a. GO enrichment of the orthogroups whose orthologous members are conserved in 7Ref-Species but lost in both *Cuscuta australis* and *Utricularia gibba*. Supplementary Data 2b. GO enrichment of the orthogroups whose orthologous members are conserved in 7Ref-Species and *Utricularia gibba*, but lost in *Cuscuta australis*. Supplementary Data 2c. GO enrichment of the orthogroups whose orthologous members are conserved in 7Ref-Species and in *Cuscuta australis*, but lost in *Utricularia gibba*. Supplementary Data 2d. GO enrichment of the principally expressed genes in *Cuscuta australis* haustoria. Supplementary Data 2e. GO enrichment of positively selected genes in *Cuscuta australis*. Supplementary Data 2f. GO enrichment of relaxed purifying selection genes in *Cuscuta australis*. Supplementary Data 2g. GO enrichment of genes in expanded gene families in *Cuscuta australis*.

File Name: Supplementary Data 3

Description: Supplementary Data 3a. Functional annotations of pseudogenes.

Supplementary Data 3b. R genes in *Cuscuta australis* and 7Ref-Species.

Supplementary Data 3c. TPS genes in *Cuscuta australis* and 7Ref-Species.

Supplementary Data 3d. P450 genes in *Cuscuta australis* and 7Ref-Species.

Supplementary Data 3e. RLK genes in *Cuscuta australis* and 7Ref-Species.

Supplementary Data 3f. Functional annotations of principally expressed genes in *Cuscuta australis* haustoria.

File Name: Supplementary Data 4

Description: Presence/absence of plastid tRNA, rRNA, and protein-coding genes in *Nicotiana tabacum*, *Ipomea nil*, four *Cuscuta spp*., and two root parasitic plants *Striga hermonthica* and *Orobanche cumana*.

File Name: Supplementary Data 5

Description: Copy numbers of Copia and Gypsy families in *Cuscuta australis* and *Ipomoea nil.*

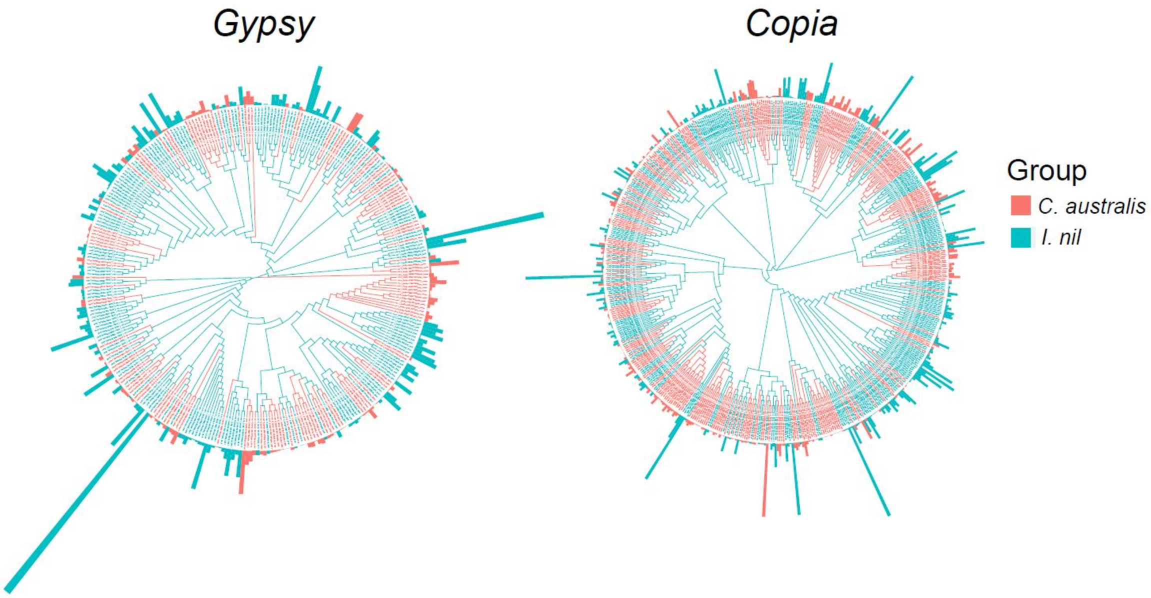


**Supplementary Figure 1. Phylogeny of *LTR/Gypsy* and *LTR/Copia* in *Cuscuta australis* and *Ipomoea nil*.** Branches of *Cuscuta australis* and *Ipomoea nil* were denoted as red and blue respectively. Bars outside dendrogram denote copy numbers of the repetitive family (detailed copy numbers can be found in Supplementary Data 5).

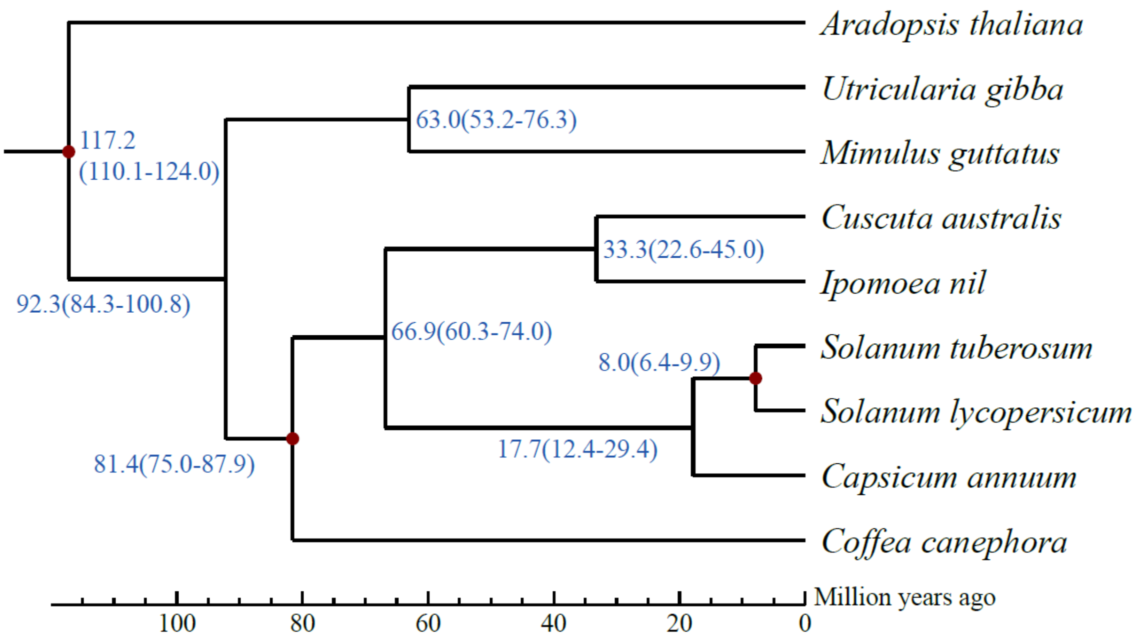


**Supplementary Figure 2. Divergence time estimation.** Divergence time was estimated using the mcmctree embedded in the PAML package. The most likely divergence times from the most recent comment ancestor are given along each node, and the estimated ranges of divergence times are shown in the parentheses. The node dots indicate the time of speciation retrieved from the Timetree database (http://timetree.org), which were used for calibrating the divergence estimation.

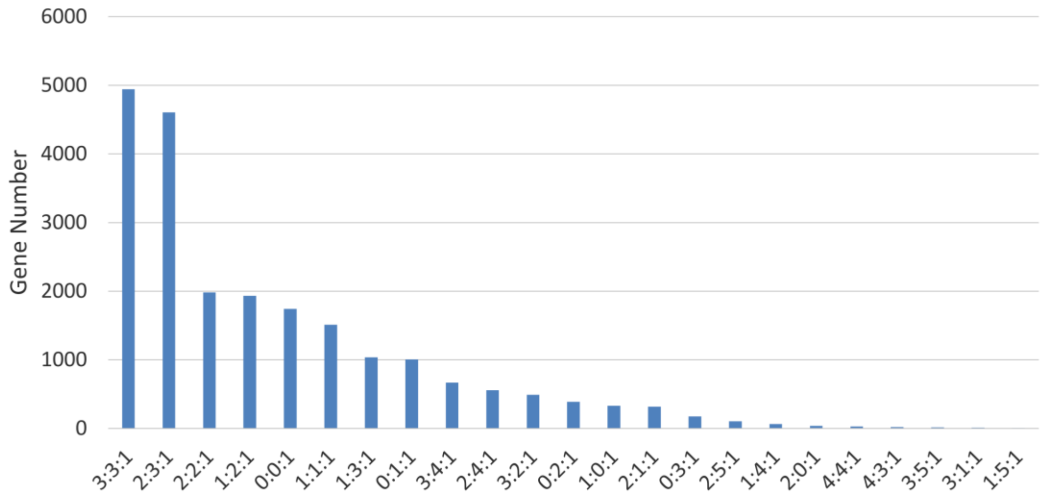


**Supplementary Figure 3. Statistics of different patterns of depths of *Cuscuta australis* and *Ipomoea nil* syntenic blocks covering all *Coffea canephora* syntenic orthologs.** Every syntenic block was anchored to the chromosomes of *Coffea canephora*, and the numbers of the syntenic blocks from *Cuscuta australis* and *Ipomoea nil* covering each individual *Coffea canephora* syntenic gene were determined to obtain the syntenic coverage information, which is described by "number of syntenic block of *Cuscuta australis*:number of syntenic block of *Ipomoea nil*:1" for each *Coffea canephora* gene locus. The syntenic *Coffea canephora* genes with a specific coverage pattern were counted and plotted.

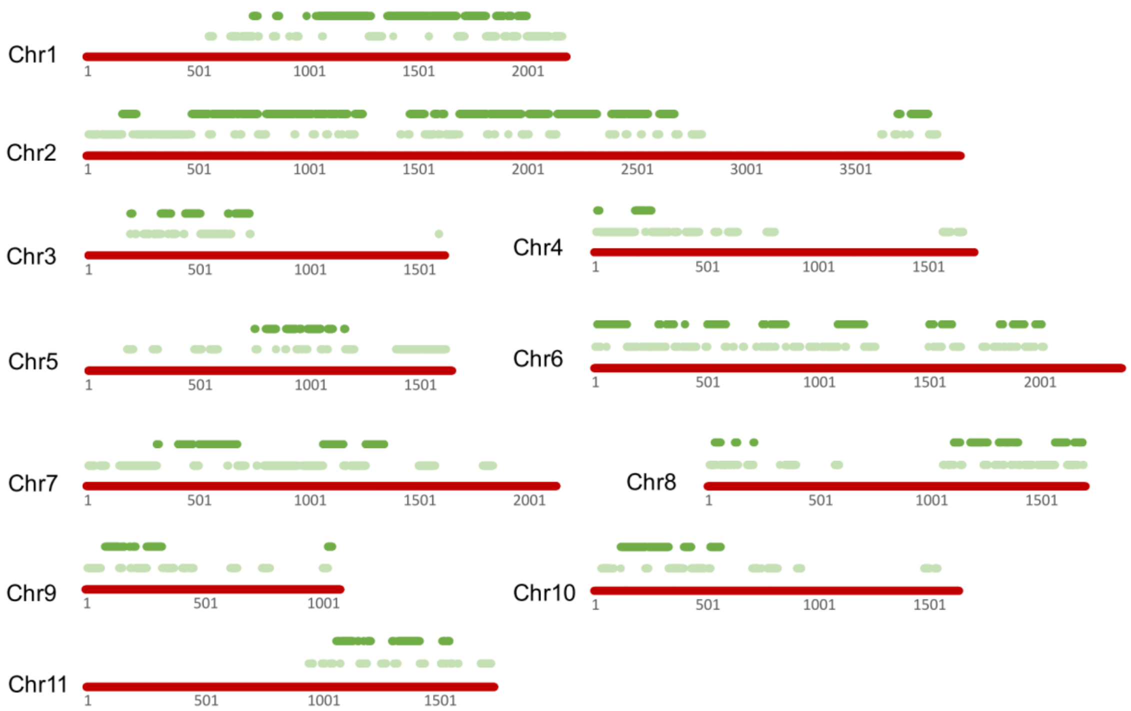


**Supplementary Figure 4. Distribution of *Coffea canephora* genes having syntenic coverages of 3:3:1 or 2:3:1 from the *Cuscuta australis* and *Ipomoea nil* syntenic blocks over the *Coffea canephora* chromosomes.** *Coffea canephora* genes having syntenic coverages 3:3:1 from *Cuscuta australis* and *Ipomoea nil* syntenic blocks are plotted as green dots, and those having 2:3:1 coverages are plotted as light green dots.

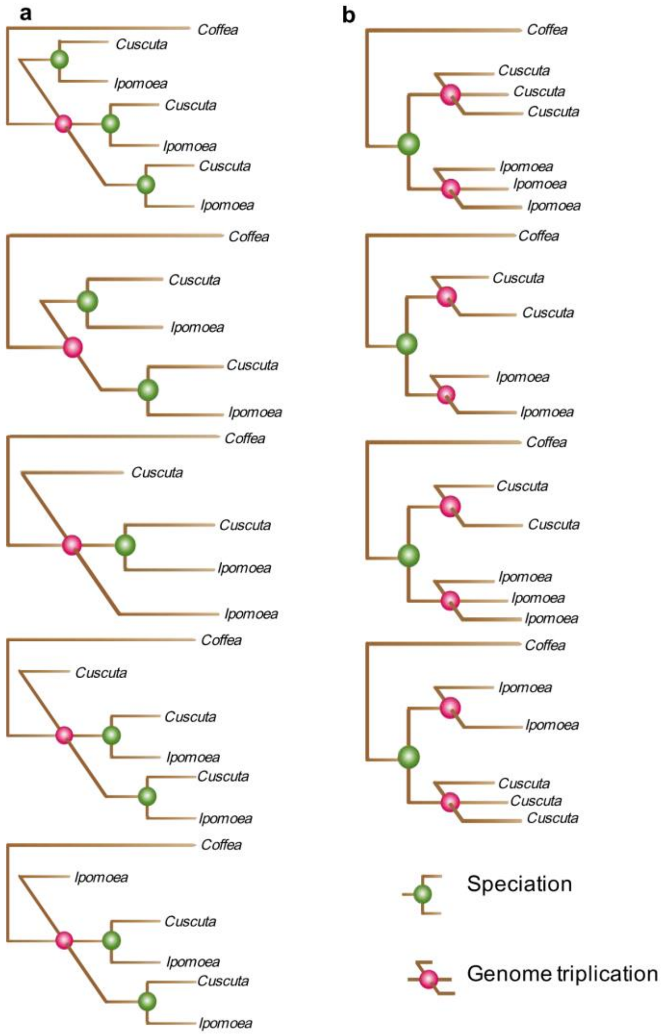


**Supplementary Figure 5. Topologies of gene trees depicting the possible scenarios of speciation among *Cuscuta australis*, *Ipomoea nil*, and *Coffea canephora*.** Orthogroups that contain one *Coffea canephora* gene, at least two *Cuscuta australis* orthologs, and at least two *Ipomoea nil* orthologs were extracted. All possible topologies were summarized after reconstructing the phylogenetic gene trees of the sequences in an orthogroup. **a.** One possible speciation scenario, in which WGD happened before the speciation between *Ipomoea* and *Cuscuta*. **b.** The alternative speciation hypothesis in which WGD happened independently after the speciation between *Ipomoea* and *Cuscuta*.

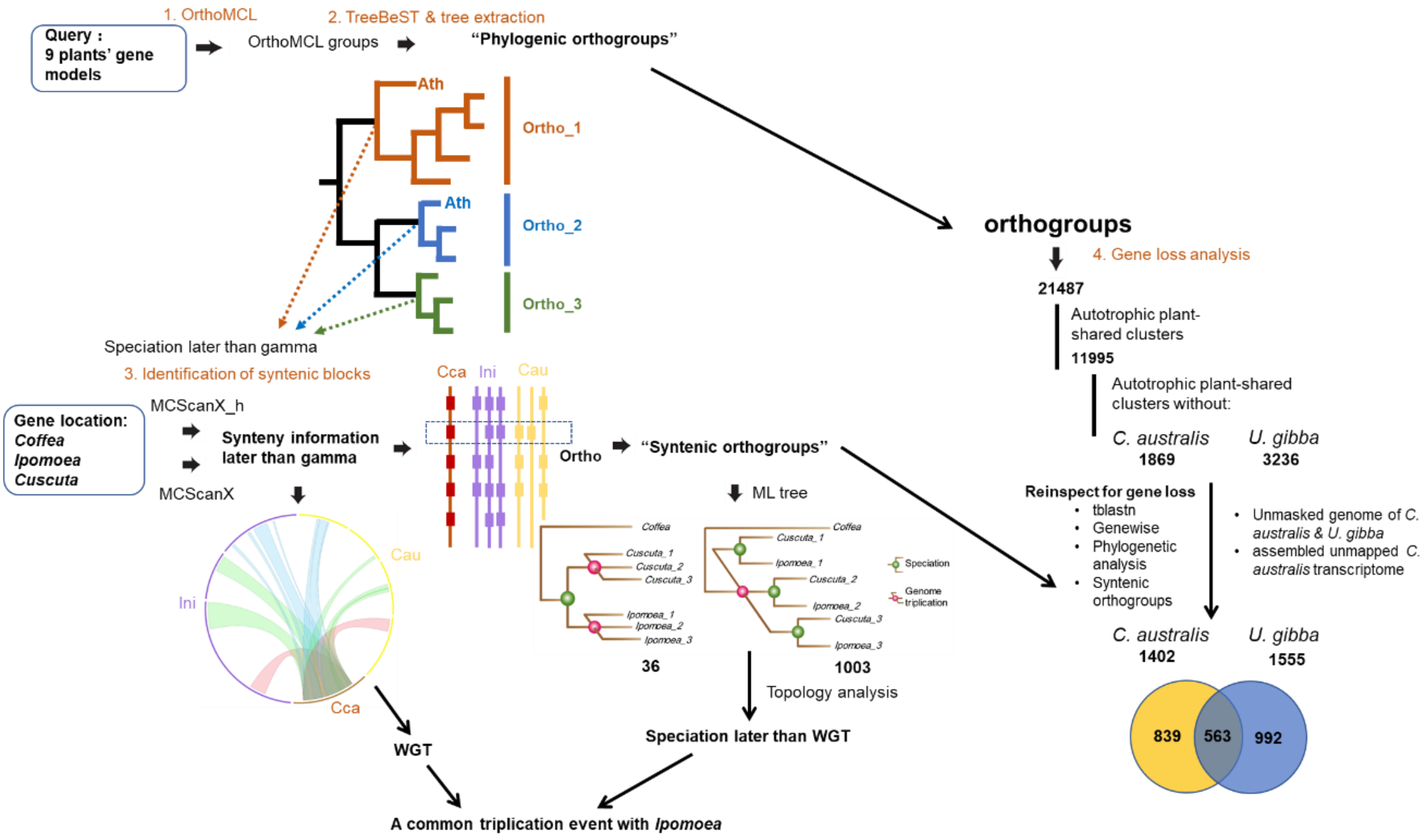
1. OrthoMCL
2. TreeBeST & tree extraction
Query :
9 plants' gene models
OrthoMCL groups
"Phylogenic orthogroups"
Ath
Ortho_1
Ath
Ortho_2
Ortho_3
Speciation later than gamma
orthogroups
4. Gene loss analysis
21487
Autotrophic plant-shared clusters
11995
Autotrophic plant-shared clusters without:
C. australis
U. gibba
1869
3236
3. Identification of syntenic blocks
Cca
Ini
Cau
Gene location:
Coffea
Ipomoea
Cuscuta
MCScanX_h
MCScanX
Synteny information later than gamma
Ortho
"Syntenic orthogroups"
ML tree
Reinspect for gene loss
tblastn
Genewise
Phylogenetic analysis
Syntenic orthogroups
Unmasked genome of C. australis & U. gibba
assembled unmapped C. australis transcriptome
Cau
Ini
Cca
Coffea
Cuscuta_1
Cuscuta_2
Cuscuta_3
Ipomoea_1
Ipomoea_2
Ipomoea_3
Speciation
Genome triplication
36
1003
C. australis
U. gibba
1402
1555
839
563
992
Topology analysis
WGT
Speciation later than WGT
A common triplication event with Ipomoea

**Supplementary Figure 6. Pipeline of orthogroup identification and syntenic analysis.** Analysis pipeline indicating the steps of identification of orthogroups and syntenic blocks. The gene models of *Cuscuta australis* and 7Ref-Species were clustered using OrthoMCL (Step 1), and resulted groups were fed to TreeBeST[1] (v1.9.2) for tree extraction (Step 2), and the "phylogenetic orthogroups" were obtained. MCScanX_h and MCScanX were employed to generate the information of the syntenic blocks (derived from speciation and gene duplication after the gamma event) in *Cuscuta australis* and the reference species (Step 3), and the information of syntenic blocks were used to obtain the "syntenic orthogroups". The "phylogenetic orthogroups" were used to analyze gene loss and "syntenic orthogroups" were used to reinspect gene loss (Step 4). The information of syntenic blocks was used for analyzing the recent WGD in *Cuscuta*, *Ipomoea*, and *Coffea*, to confirm whether whole-genome triplication (WGT) happened during the evolution of *Cuscuta* and *Ipomoea*. The phylogenetic relationship of the orthologs among *Cuscuta australis*, *Ipomoea nil*, and *Coffea canephora* were examined to conclude the order of WGT and speciation between *Cuscuta* and *Ipomoea*. had a common whole-genome triplication was concluded. Based on these lines of evidence, that *Cuscuta* and *Ipomoea* had a common WGT event before the split from their common ancestor was concluded.

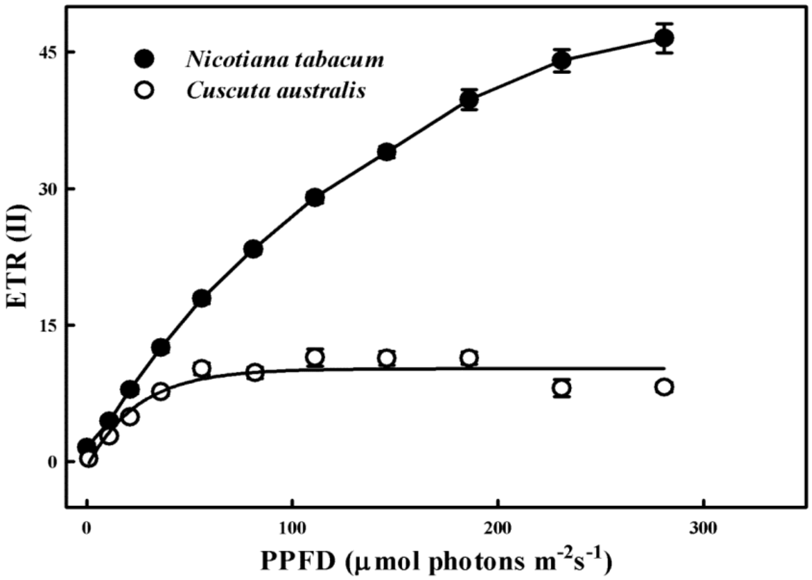


**Supplementary Figure 7. Light-response changes in photosynthetic electron flow through PSII (ETR(II)) in *Cuscuta australis* stems and leaves of *Nicotiana tabacum*.**

Data are means ± SE (n = 5).

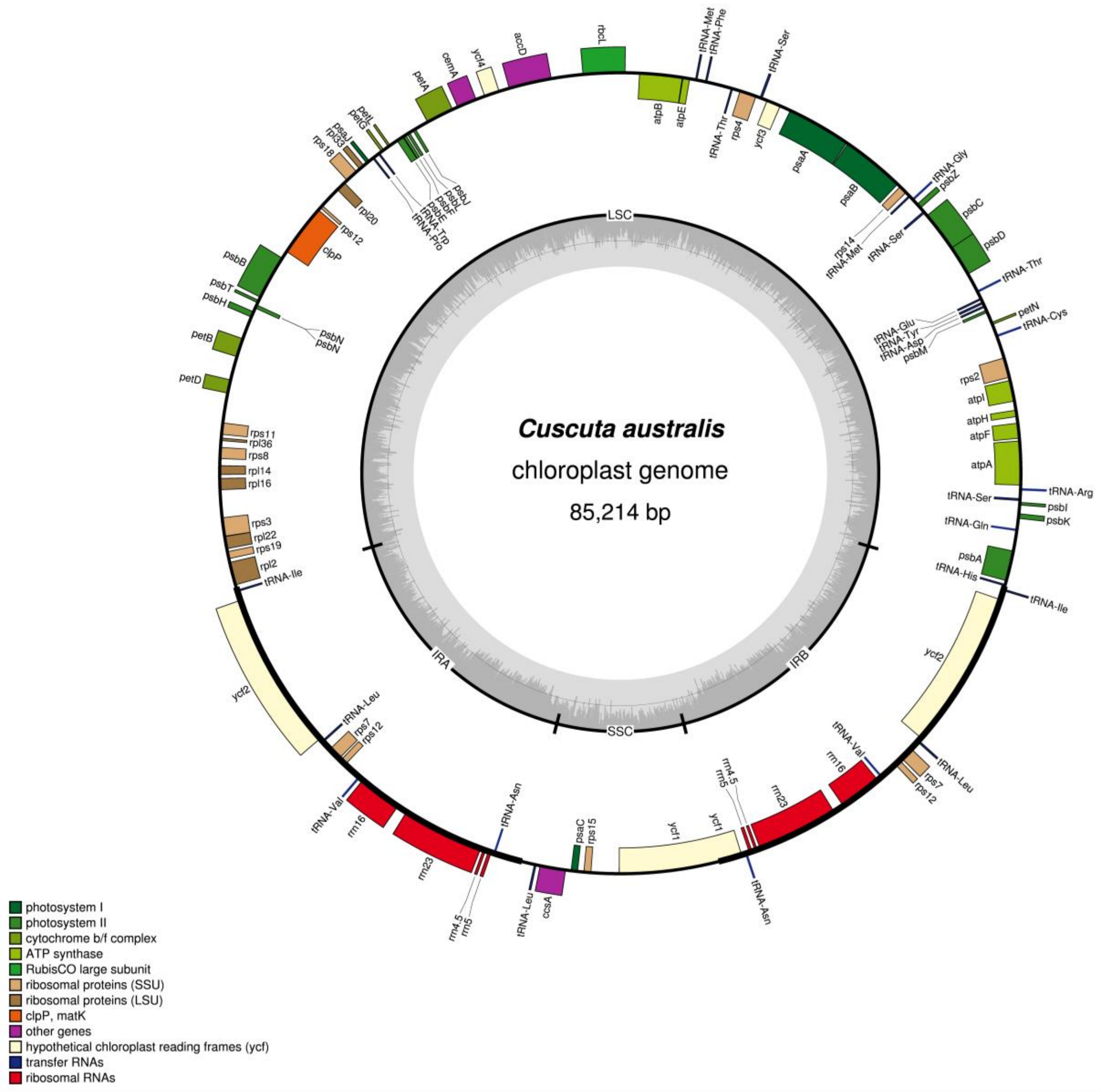


**Supplementary Figure 8. Circular map of the complete plastid genome of *Cuscuta australis.***

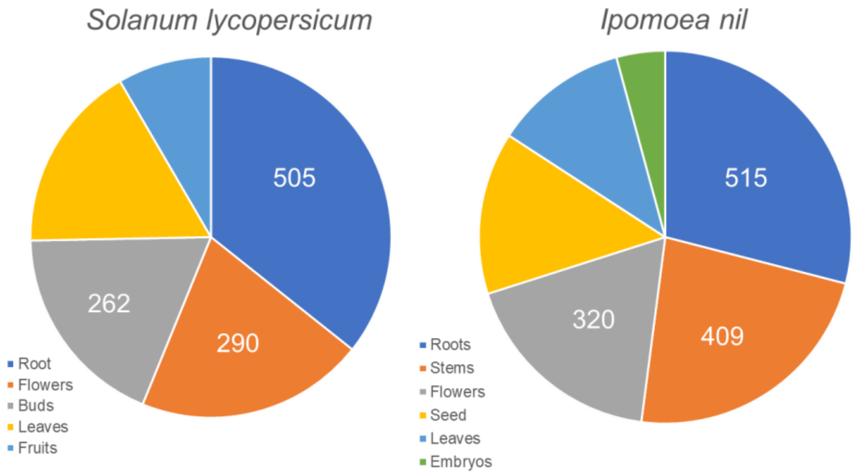


**Supplementary Figure 9. The principally expressed tissues (PETs) of *Cuscuta. australis* haustorium principally expressed genes' orthologues in *Solanum lycopersicum* and *Ipomoea. nil*.** The respective PETs of the orthogroups, which have principally expressed genes in *C. australis* haustoria were identified in *Solanum lycopersicum* and *Ipomoea. nil*. The numbers of orthogroups, which have PETs in leaves, roots, flowers, and other tissues, are shown in the respective slices of the pie charts. Only the slices of the top three PETs are shown with numbers.

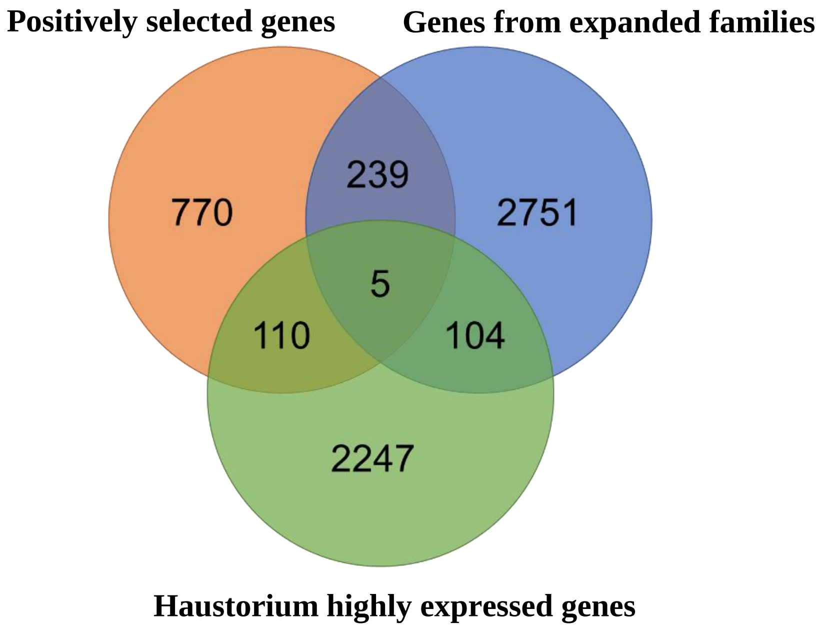


**Supplementary Figure 10. Venn diagram of the positively selected genes, genes from the expanded gene families, and genes highly expressed in prehaustoria/haustoria**.

**Supplementary Table 1. Genome assembly statistics of *Cuscuta australis***

| Type | Contigs | | Scaffolds | | Superscaffolds | |
|---|---|---|---|---|---|---|
| | **Size** | **Number** | **Size** | **Number** | **Size** | **Number** |
| N90 | 904385 | 87 | 1850081 | 46 | 1850081 | 46 |
| N80 | 1418971 | 64 | 2904980 | 34 | 2904980 | 34 |
| N70 | 2100488 | 48 | 3950442 | 27 | 3950442 | 27 |
| N60 | 2542181 | 36 | 4610076 | 21 | 4610076 | 21 |
| N50 | 3625894 | 27 | 5945234 | 16 | 5945234 | 16 |
| N40 | 4249895 | 20 | 7261444 | 12 | 7261444 | 12 |
| N30 | 4403227 | 14 | 7763232 | 8 | 7763232 | 8 |
| N20 | 4855865 | 8 | 10192992 | 5 | 10192992 | 5 |
| N10 | 6540893 | 4 | 10743034 | 3 | 10743034 | 3 |
| Total Length | 267146983 | 255 | 267213323 | 103 | 267213323 | 103 |
| Maximum Length | 10192992 | - | 13922953 | | 13922953 | |
| Minimum Length | 5448 | - | 10000 | | 10000 | |

**Supplementary Table 2. Contigs containing organellar sequences**

| | Contig_ID | Length | Average depth | Note |
|---|---|---|---|---|
| Plastid | C141C | 85214 | 2593.871433 | complete plastome |
| | C131M | 186399 | 1214.601551 | |
| | C135M | 146508 | 1685.295412 | |
| | C136M | 141688 | 1072.910925 | |
| | C139M | 134523 | 1862.792701 | |
| | C140M | 123029 | 1711.995977 | |
| | C143M | 97161 | 1609.256254 | |
| | C146M | 87099 | 1842.503 | |
| | C149M | 81803 | 1723.384704 | |
| | C154M | 77475 | 1336.538172 | |
| | C162M | 57724 | 1447.579904 | |
| | C173M | 50918 | 1344.030801 | |
| | C175M | 50434 | 778.2785238 | |
| Mitochondria | C176M | 50271 | 1563.827082 | |
| | C185M | 48206 | 1842.295907 | |
| | C186M | 45851 | 1500.950283 | |
| | C189M | 45272 | 928.8866392 | |
| | C192M | 43229 | 1163.849815 | |
| | C193M | 43010 | 1248.797399 | |
| | C196M | 41178 | 1140.851044 | |
| | C202M | 39887 | 1245.670002 | |
| | C208M | 36278 | 1340.751504 | |
| | C220M | 33415 | 1163.144485 | |
| | C221M | 33114 | 1031.367691 | |
| | C222M | 32823 | 962.1748523 | |
| | C225M | 29664 | 1132.485868 | |

| | | |
|---|---|---|
| C235M | 22150 | 582.2652773 |
| C238M | 18912 | 1009.900755 |
| C246M | 14731 | 1096.597367 |
| C253M | 8634 | 832.8695944 |

**Supplementary Table 3. Accuracy and heterozygosity of *Cuscuta australis* genome assembly**

| **Errors in Pacbio assembly before Pilon correction** | |
|---|---|
| Numbers of homozygous SNPs | 24636 |
| Numbers of homozygous INDELs | 115496 |
| Estimated accuracy | 0.9995 |
| **Errors in Pacbio assembly after Pilon correction** | |
| Numbers of homozygous SNPs | 23656 |
| Numbers of homozygous INDELs | 13671 |
| Estimated accuracy | 0.9999 |
| **Within genome heterozygosity** | |
| Within genome heterozygous SNPs | 29070 |
| Numbers of heterozygous INDELs | 4733 |
| Estimated within genome heterozygosity | 0.013% |

**Supplementary Table 4. Repetitive elements in *Cuscuta australis* genome**

| | ***Ipomoea nil*** | | | ***Cuscuta australis*** | | |
|---|---|---|---|---|---|---|
| **Classes** | **Copies** | **Total size** | **Percentage in genome** | **Copies** | **Total size** | **Percentage in genome** |
| DNA Transposon | 237,062 | 79,245,270 | 10.8% | 73,415 | 19,401,328 | 7.3% |
| LTR | 191,314 | 172,365,468 | 23.5% | 79,011 | 63,746,515 | 23.9% |
| LINE | 28,561 | 19,757,447 | 2.7% | 18,039 | 6,456,063 | 2.4% |
| SINE | 25,188 | 6,566,780 | 0.9% | 8,131 | 1,239,886 | 0.5% |
| Helitron | 7,778 | 4,266,513 | 0.6% | 7,138 | 2,852,074 | 1.1% |
| Simple Repeats | 366,864 | 22,582,319 | 3.1% | 85,340 | 6,672,582 | 2.5% |
| Unknown | 623,410 | 181,060,611 | 24.6% | 259,895 | 62,796,207 | 23.5% |
| Total | - | 474,203,812 | 64.5% | - | 155,040,159 | 58% |

**Supplementary Table 5. Two cluster analyses of *Cuscuta australis* and related species.**

| Outgroup | Ingroup A | Ingroup B | bA | bB | delta | s.e. | Z | CP | Faster |
|---|---|---|---|---|---|---|---|---|---|
| *Arabidopsis thaliana* | *Cuscuta australis* | *Solanum tuberosum* | 0.206047 | 0.179943 | 0.026103 | 0.001004 | 25.989989 | 99.96% | *Cuscuta australis* |
| *Arabidopsis thaliana* | *Cuscuta australis* | *Solanum lycopersicum* | 0.20728 | 0.170191 | 0.037088 | 0.000958 | 38.707126 | 99.96% | *Cuscuta australis* |
| *Arabidopsis thaliana* | *Cuscuta australis* | *Coffea canephora* | 0.2209 | 0.185532 | 0.035368 | 0.000985 | 35.891519 | 99.96% | *Cuscuta australis* |
| *Arabidopsis thaliana* | *Cuscuta australis* | *Capsicum annuum* | 0.207216 | 0.177334 | 0.029883 | 0.000985 | 30.338771 | 99.96% | *Cuscuta australis* |
| *Arabidopsis thaliana* | *Cuscuta australis* | *Ipomoea nil* | 0.125598 | 0.094319 | 0.031279 | 0.000767 | 40.798868 | 99.96% | *Cuscuta australis* |
| *Arabidopsis thaliana* | *Cuscuta australis* | *Mimulus guttatus* | 0.225954 | 0.211697 | 0.014257 | 0.001028 | 13.865957 | 99.96% | *Cuscuta australis* |

The branch length was calculated using *Arabidopsis thaliana* as the outgroup. Z-statistic was used to test whether the distances between ingroups (bA, bB) to the outgroup is significantly different from 0 or not. Delta is the absolute difference between bA and bB. Z-statistics (Z) is delta/standard error (s.e.). CP (confident probability) is equal to 1 - (p value).

**Supplementary Table 6. Relative rate test of *Cuscuta australis* and related species**

The chi-square test was based on 1 degree of freedom.

| Outgroup | Ingroup 1 | Ingroup 2 | Identical | Ingroup 1 specific | Ingroup 2 specific | Chi-score | p-value | Faster |
|---|---|---|---|---|---|---|---|---|
| *Arabidopsis thaliana* | *Cuscuta australis* | *Solanum tuberosum* | 467586 | 100123 | 88818 | 676.42 | <0.00001 | *Cuscuta australis* |
| *Arabidopsis thaliana* | *Cuscuta australis* | *Solanum lycopersicum* | 504389 | 107675 | 90153 | 1551.96 | <0.00001 | *Cuscuta australis* |
| *Arabidopsis thaliana* | *Cuscuta australis* | *Coffea canephora* | 498947 | 113425 | 96542 | 1357.53 | <0.00001 | *Cuscuta australis* |
| *Arabidopsis thaliana* | *Cuscuta australis* | *Capsicum annuum* | 484351 | 103773 | 90260 | 941.08 | <0.00001 | *Cuscuta australis* |
| *Arabidopsis thaliana* | *Cuscuta australis* | *Ipomoea nil* | 537711 | 69320 | 54480 | 1778.88 | <0.00001 | *Cuscuta australis* |
| *Arabidopsis thaliana* | *Cuscuta australis* | *Mimulus guttatus* | 485175 | 113596 | 106831 | 207.62 | <0.00001 | *Cuscuta australis* |

**Supplementary Table 7. Statistics of different patterns of depths of *Cuscuta australis* and *Ipomoea nil* syntenic blocks covering all *Coffea canephora* syntenic orthologs**

| *Cuscuta australis*:*Ipomoea nil*: *Coffea canephora* | Count |
|---|---|
| 3:3:1 | 4943 |
| 2:3:1 | 4604 |
| 2:2:1 | 1981 |
| 1:2:1 | 1931 |
| 0:0:1 | 1740 |
| 1:1:1 | 1513 |
| 1:3:1 | 1037 |
| 0:1:1 | 1007 |
| 3:4:1 | 668 |
| 2:4:1 | 557 |
| 3:2:1 | 490 |
| 0:2:1 | 390 |
| 1:0:1 | 330 |
| 2:1:1 | 316 |
| 0:3:1 | 174 |
| 2:5:1 | 104 |
| 1:4:1 | 66 |
| 2:0:1 | 39 |
| 4:4:1 | 28 |
| 4:3:1 | 19 |
| 3:5:1 | 16 |
| 3:1:1 | 13 |
| 1:5:1 | 3 |

Note: For this statistic, every syntenic block was anchored to the chromosomes of *Coffea canephora*, and the numbers of the syntenic blocks from *Cuscuta australis* and *Ipomoea nil* covering each individual *Coffea canephora* syntenic gene were determined to obtain the syntenic coverage information, which is described by "number of syntenic block of *Cuscuta australis*:number of syntenic block of *Ipomoea nil*:1" for each *Coffea canephora* gene locus. The syntenic *Coffea canephora* genes with a specific coverage pattern were counted and plotted.

**Supplementary Table 8. Mapping ratios of RNA-Seq reads from *Cuscuta australis* and *Cuscuta pentagona* to the PacBio assembly of *C. australis***

| Sample Name | Species Name | Sample ID | Hosts | Contamination Percentage | Alignment Ratio |
|---|---|---|---|---|---|
| **C10_Seeds** | *Cuscuta australis* | C10 | - | - | 96.06% |
| **C1_Bud on soybean host** | *Cuscuta australis* | C1 | Soybean | 0.34% | 96.40% |
| **C3_Ovary on soybean host** | *Cuscuta australis* | C3 | Soybean | 0.34% | 96.87% |
| **C4_Germinating seeds** | *Cuscuta australis* | C4 | - | - | 95.28% |
| **C5_Haustoria on tomato host** | *Cuscuta australis* | C5 | Tomato | 0.44% | 96.75% |
| **C6_Prehaustoria on tomato host** | *Cuscuta australis* | C6 | Tomato | 3.35% | 96.39% |
| **C7_Curling stems on tomato host** | *Cuscuta australis* | C7 | Tomato | 0.13% | 96.87% |
| **C8_Stem tip on tomato host** | *Cuscuta australis* | C8 | Tomato | 1.21% | 96.83% |
| **flowers_tobacco*** | *Cuscuta pentagona* | flowers_tobacco.rep1 | Tobacco | 2.36% | 78.38% |
| | | flowers_tobacco.rep2 | Tobacco | 4.53% | 79.30% |
| | | flowers_tobacco.rep3 | Tobacco | 2.65% | 80.55% |
| | | flowers_tobacco.rep4 | Tobacco | 1.81% | 77.39% |
| **flowers_tomato*** | *Cuscuta pentagona* | flowers_tomato.rep1 | Tomato | 3.08% | 79.93% |
| | | flowers_tomato.rep2 | Tomato | 9.91% | 75.31% |
| | | flowers_tomato.rep3 | Tomato | 3.51% | 78.42% |
| | | flowers_tomato.rep4 | Tomato | 9.94% | 75.32% |
| **haustorial_stems_tobacco*** | *Cuscuta pentagona* | haustorial_stems_tobacco.rep1 | Tobacco | 3.45% | 61.17% |
| | | haustorial_stems_tobacco.rep2 | Tobacco | 4.33% | 61.09% |
| | | haustorial_stems_tobacco.rep3 | Tobacco | 2.22% | 68.85% |
| | | haustorial_stems_tobacco.rep4 | Tobacco | 3.04% | 68.86% |
| **haustorial_stems_tomato*** | *Cuscuta pentagona* | haustorial_stems_tomato.rep1 | Tomato | 1.60% | 63.80% |
| | | haustorial_stems_tomato.rep2 | Tomato | 2.29% | 71.41% |
| | | haustorial_stems_tomato.rep3 | Tomato | 2.95% | 64.19% |
| | | haustorial_stems_tomato.rep4 | Tomato | 2.13% | 71.30% |

| | | | | | |
|---|---|---|---|---|---|
| **prehaustorial_stems_tobacco*** | *Cuscuta pentagona* | prehaustorial_stems_tobacco.rep1 | Tobacco | 4.94% | 78.25% |
| | | prehaustorial_stems_tobacco.rep2 | Tobacco | 2.77% | 79.56% |
| | | prehaustorial_stems_tobacco.rep3 | Tobacco | 2.59% | 80.34% |
| | | prehaustorial_stems_tobacco.rep4 | Tobacco | 2.94% | 76.26% |
| **prehaustorial_stems_tomato*** | *Cuscuta pentagona* | prehaustorial_stems_tomato.rep1 | Tomato | 2.20% | 72.39% |
| | | prehaustorial_stems_tomato.rep2 | Tomato | 3.09% | 76.03% |
| | | prehaustorial_stems_tomato.rep3 | Tomato | 2.06% | 74.68% |
| | | prehaustorial_stems_tomato.rep4 | Tomato | 1.54% | 77.65% |
| **seedlings*** | *Cuscuta pentagona* | seedlings.rep1 | - | | 80.23% |
| | | seedlings.rep2 | - | | 79.47% |
| | | seedlings.rep3 | - | | 80.97% |
| | | seedlings.rep4 | - | | 79.88% |
| **seeds*** | *Cuscuta pentagona* | seeds.rep1 | - | | 72.01% |
| | | seeds.rep2 | - | | 71.94% |
| | | seeds.rep3 | - | | 67.84% |
| | | seeds.rep4 | - | | 71.80% |
| **stems_tobacco*** | *Cuscuta pentagona* | stems_tobacco.rep1 | Tobacco | 6.10% | 80.70% |
| | | stems_tobacco.rep2 | Tobacco | 1.94% | 80.75% |
| | | stems_tobacco.rep3 | Tobacco | 6.13% | 80.76% |
| | | stems_tobacco.rep4 | Tobacco | 4.88% | 75.67% |
| **stems_tomato*** | *Cuscuta pentagona* | stems_tomato.rep1 | Tomato | 4.33% | 79.29% |
| | | stems_tomato.rep2 | Tomato | 4.07% | 80.58% |
| | | stems_tomato.rep3 | Tomato | 3.71% | 80.08% |
| | | stems_tomato.rep4 | Tomato | 3.91% | 80.51% |

*RNA-Seq data of *Cuscuta pentagon* was obtained from Reference [2]. Note: Reads of host mRNAs have been filtered out before mapping was done.

### Supplementary Notes

#### Supplementary Note 1.

##### Phylogeny and divergence times.

Phylogenetic tree was build using RAxML[3] (v8.2.11) with the protein sequences of single-copy orthogroups (Fig. 1b). A long branch was detected in *Cuscuta australis*, suggesting a rapid evolution rate. This was further validated using two cluster analysis and Tajima's relative rate test[4-6] (Supplementary Table 5 and Supplementary Table 6). Divergence time was estimated using the mcmctree (burn-in=10,000, sample-number=100,000, sample-frequency=2) embedded in the PAML package[7] (v4.9e) with calibration set to 106.8-124.8 Mya between *Arabidopsis thaliana* and *Mimulus guttatus* and 6.5-7.4 Mya between *Solanum lycopersicum* and *Solanum tuberosum* (Supplementary Fig. 2). These divergence times were obtained from timetree (http://www.timetree.org).

#### Supplementary Note 2.

##### Expansion and contraction of *Cuscuta australis* gene families.

The protein sequences from *Cuscuta australis*, *Utricularia gibba*, and the 7Ref-Species were firstly all-vs-all aligned using blastp[8] (v2.4.0). E-value of each hit was transformed into *W* index [1] ranging from 0 to 100 using the following formula.

$$W = Min(100, \mathrm{ROUND}(\frac{-\log_{10}(Evalue)}{2}))$$

Thereafter, *W* indices were passed to hcluster[1] (v0.5.1) as weight matrices to build gene families with the parameters "-m 170 -w 0 -s 0.34 -O". A total of 13981 gene families were constructed using this method, covering 92% of the total. To directly compare sizes of gene families between species, we computed the *F*-indices, which describe the size differences of conserved gene families in 7Ref-Species (conserved gene families must exist in *Arabidopsis* and at least five out of the six remaining species in the 7Ref-Species), using the formula below.

In this formula, *i* refers to a given species and *j* refers to a given family, $c_{ij}$ is the number of genes in *j* family in *i* species, and $N_j$ is the total number of genes in *j* family of all species, and *S* represents the number of species. *F*-index is a modified log2-fold-change coefficient ($\log_2 \frac{c_{ij}}{N_j}$), so the relationship between *F*-index and $\log_2 \frac{c_{ij}}{N_j}$ is linear. *F*-indices range from 0 to 1, and when $F_{ij} = 0.5$, in species *i*, the gene number in gene family *j* equals to the average size of this gene family in all species; if $F_{ij} = 0$, there are no genes in family *j* in species *i*, and if $F_{ij} = 1$, it indicates that only species *i*, but not the others, harbors the gene family *j*. Thereafter, a box plot of *F*-indices of the families (data are listed in Supplementary Data 1a), that are conserved in 7Ref-Species, for all species was constructed (Fig. 2b).

$$F_{ij} = \frac{\log_2\left(a\frac{c_{ij}}{N_j} + \frac{1}{2}\right) + 1}{\log_2\left(a + \frac{1}{2}\right) + 1}$$

$$a = \frac{S^2 - 2S}{2}$$

To investigate expansions and contractions of gene families, we used the gain-and-death (GD) model in BadiRates[9] (v1.35), a program that estimates family turnover rates by likelihood-based methods. Species tree and gene family number in each species were used as the input file for BadiRates[9]. To calculate which lineages were significantly expanded or contracted, we used the free model in BadiRates to estimate the sizes of the ancestral gene families in all clades of the species tree. For the branches whose gene families did not experience family size changes, they were set to be the background branches, which have the same family turnover rate; thereafter, based on this model, we re-estimated the likelihood, and the model was regarded as the null hypothesis. For branches that experienced size changes, an alternative hypothesis for each branch was built by forcing the given branch to follow the same turnover rate with the background branches. A branch that experienced size changes was considered to be significant, if AIC (alternative hypothesis) - AIC (null hypothesis) > 2 (Akaike's information criterion (AIC)[10] was computed from the likelihood and numbers of parameters in each model). Hereafter, the significantly expanded and contracted gene families in all eight species were obtained (details are in Supplementary Data 1a).

**Supplementary Note 3.**

**Identification of gene loss in the *Cuscuta australis* and *Utricularia gibba* genome**

We developed a pipeline to identify the orthologous gene clusters in the genomes of *Cuscuta australis, Utricularia gibba*, and the 7Ref-Species (Supplementary Fig. 6), which considers the phylogenetic and the collinear information of genes. In order to obtain a rather complete overview of the lost genes, pseudogenes in *Cuscuta australis* and *Utricularia gibba* were also integrated into the gene models, by masking the frame-shift and premature stop codon mutation sites to X and N on the amino acid and nucleotide level, respectively. Considering the possibility that genes with frame-shifts and premature stop codons identified *in silico* may still be translated to proteins which retain their functions, these genes were treated as normal genes and input into the gene loss analysis.

The details of the pipeline procedure are summarized as the following:

**1. Identification of the orthogroups from phylogenetic trees**

All the predicted gene models in the nine species were clustered using orthoMCL[11] (v2.0.9) with the parameters set as percentMatchCutoff=50, evalueExponentCutoff=-5, inflation=1.5, resulting in 25,330 OrthoMCL groups; phylogenetic analysis was carried out for each group obtained above using protein alignments produced by t-Coffee[12] (v11.00) (aligner set: mafftgins_msa, muscle_msa, kalign_msa, and t_coffee_msa), and the codon alignments were deduced from protein alignments. To further optimize the tree topologies, we used TreeBeST[1] (v1.9.2) to construct five different types of phylogenetic trees: two maximum likelihood (ML) trees based on protein alignments and codon alignments, respectively, and three neighbour-joining (NJ) trees based on codon alignments with p-distances, dN distances, and dS distances. The final trees were generated by the TreeBeST-implemented tree merging algorithm, which integrates species tree and the five trees, considering the advantages of DNA- and protein-based trees, bootstrap values, and the minimization of gene duplication/loss numbers.

*Arabidopsis* is the outgroup for *Cuscuta*, *Utricularia*, and the other six autotrophic reference species belong to the lamiids; thus, all the genes in each orthogroup must have the same *Arabidopsis* ortholog; thus, these genes in a specific orthogroups should closely cluster with the *Arabidopsis* outgroup ortholog gene in a branch of the gene family phylogenetic tree. Since *Arabidopsis* is the most remotely related with the other species and has been intensively studied for gene functions, tree topologies can be constructed unambiguously and the functions of orthogroups can be assigned accordingly. To divide each phylogenetic tree into individual orthogroups, we firstly chose the clades containing *Arabidopsis* outgroups and then treated the remaining clades with the same levels as those containing *Arabidopsis* outgroups as independent orthogroups. A total of 21,487 orthogroups (≥ 2 species) were identified using this method, among which, 11,995 are conserved in the 7Ref-Species (considering that there could be mistakes in the genome annotation of 7-Ref species, a "conserved orthogroup" is formed only if it contains the orthologs from *Arabidopsis* and all or five out of the remaining six species from 7Ref-Species), including 16,799 *Arabidopsis* genes (Supplementary Data 1b and 1c). Among the conserved orthogroups, 1869 and 3236 orthogroups in *Cuscuta australis* and *Utricularia gibba*, respectively, were identified to potentially have gene losses (Supplementary Data 1b). These orthogroups are named as orthogroups with potential gene losses (OPGL).

Various factors could affect gene loss identification, including OrthoMCL grouping, phylogenetic reconstruction of gene trees, and genome masking and incomplete transcriptome information during the annotation step. To minimize false positives that could be caused by these factors, the previously identified gene losses in both *Cuscuta australis* and *Utricularia gibba* were further inspected by the following pipeline (Supplemental Fig. 6):

1) tblastn (E-value threshold = 1e-10) search was first applied to search the unmasked *Cuscuta australis* genome and the assembled unmapped *Cuscuta australis* transcriptome (See methods "Transcriptome analysis" for detail) with all the genes in the 1869 orthogroups, which appeared to have lost *Cuscuta australis* members.

After tblastn search of the *Cuscuta australis* genome, the resulted hits were concatenated (because of possible introns), if the distances of the adjacent matches were within 10 kb. In this manner, we could find all the homologs of the genes in all OPGLs in *Cuscuta australis*, if there were any. We performed the same analysis for *Utricularia gibba.*

2) All the homologs obtained from the Step a. were aligned against the annotated genes in the respective *Cuscuta australis* or *Utricularia gibba* genome. If a homolog had been annotated (covering at least 80%), it was regarded as successfully annotated; otherwise, the homolog was assigned as unannotated.

3) The sequences of all the newly identified unannotated homologs were extended to include 5 kb of both upstream and downstream border sequences. Genewise[13] (v2.2.0) was exploited to annotate the gene models of these homologs, and the query sequences used in tblastn search were used as the Genewise references. The output protein sequences from Genewise annotation must cover at least 50% of the reference sequences. Since multiple reference sequences from different genomes were used as queries, redundant output protein sequences exist. The redundant sequences were removed with CD-HIT[14] (v4.6) (identity threshold 90%), leaving only the longest sequences.

4) All the sequences obtained from Step c. and the gene models from 7-Ref species and the previously annotated *Cuscuta australis* and *Utricularia gibba* were integrated into a database. All the sequences in each OPGL were used as queries to perform blastp[8] (v2.4.0) (E-value threshold 1e-5) against this database. The resulted subjects from each query in a given OPGL were scored by transforming the E-value of each

subject into *W* index (See Supplementary Note 2. "Construction of gene families"). After all the genes in a given OPGL are done with blastp as queries, the scores of the same subjects were summed to obtain their final scores. Then, the subjects identical to those in the OPGL were removed from further analysis. The remaining subjects were subsequently classified according their species, and the top five subjects with the highest scores were acquired for each species.

5) These subjects and the sequences in the respective OPGL were used for phylogenetic analysis (clustalW and Fasttree; reliable clades must have bootstrap values greater than 80%). If certain genes from *Cuscuta australis* or *Utricularia gibba* appeared in the clade of OPGL, these orthogroups are no longer considered to be involved in gene loss.

A total of 1494 and 1555 orthogroups were still identified to have gene loss in *Cuscuta australis* and *Utricularia gibba*, respectively.

**2. Identification of orthogroups from collinear fragment information**

Although the method described in "1. Identification of the orthogroups from phylogenetic trees" can generate almost complete information of orthogroups using protein sequences, it ignores the synteny information among different species. Thus, a pipeline was developed to identify orthogroups based on syntenic gene pairs (Supplementary Fig. 6), in order to complement the orthogroups obtained from the above step.

Firstly, protein sequences from *Cuscuta australis*, *Arabidopsis thaliana*, *Coffea canephora*, and *Ipomoea nil* were all-vs-all aligned using blastp[8] (v2.4.0) with the E-value set to 1e-5. Then the results from blastp and the information of gene loci in all *Cuscuta australis*, *Arabidopsis thaliana*, *Coffea canephora*, and *Ipomoea nil* were used to feed MCScanX[15] (version 201610) to obtain the information about the collinear fragments between all pairs of these four species.

These fragments were originated from two processes, namely, genome duplication (including the gamma triplication event, before the speciation between lamiids and Arabidopsis) and the divergence between *Cuscuta australis*, *Arabidopsis thaliana*, *Coffea canephora*, and *Ipomoea nil*. To construct orthogroups, we utilized the previously established orthogroups (from “1) Identification of the orthogroups from phylogenetic trees”) to identify a set of collinear fragments derived from speciation (MCScanX_h (version 201610)). These fragments were further used as a reference to guide the selection of MCScanX results, resulting in fragments that are suitable for identification of orthology (the noise of paralogs generated by genome duplication could be removed).

In this manner, 54,165 genes among the 116,404 genes in *Cuscuta australis*, *Arabidopsis thaliana*, *Coffea canephora*, and *Ipomoea nil* were clustered into 14,978 orthogroups. Using these orthogroups based on collinearity, the orthogroups obtained from phylogenetic analysis were corrected: 93 out of 1495 orthogroups, which were formerly considered to have no *Cuscuta australis* members, could retrieve their *Cuscuta australis* orthologs.

Functions of all conserved orthogroups were assigned based on the *Arabidopsis* orthologs (Supplementary Data 1b and 1c), and GO enrichment of the these 1402 orthogroups whose *Cuscuta australis* members are absent indicated that “transport”, “photosynthetic”, “regulation of transcription”, and “response to stimulus” were overrepresented (Supplementary Data 2a, Supplementary Data 2b).

**3. Manual inspection on the lost genes of interest.**

To manually inspect the lost genes in *Cuscuta australis*, which are mentioned in the main text and whose orthologs in *Arabidopsis* play important roles in leaf and root development, nutrient transport, flowering, and defenses, etc. (Supplementary Data 1c), the following procedure was adopted:

1) Inspection of the PacBio assembly

a Exonerate[16] (v2.2.0) software was used (--model protein2genome --percent 50) to map the protein sequences of *Arabidopsis* orthologs to *Cuscuta australis* PacBio assembly, in order to ensure the target gene loci have been annotated as functional or pseudogenes.

b Each *Arabidopsis* protein sequence was set as a query to perform blastp[8] (v2.4.0) against the gene models of *Cuscuta australis* and 7Ref-Species (E-value=1e-5). The obtained subject sequences were used to conduct multiple-sequence alignment (MSA) with the MUSCLE[17] (v3.8.31) software (in case of too many subject sequences, i.e. over 100, MCL (inflation=1.4) was used to cluster the subject sequences into smaller groups, and the group containing the query was used for MSA). The sequences with poor alignment quality in MSA were manually removed to redo MSA; this procedure was repeated until no sequences were considered to have poor alignment quality. Fasttree[18] (v2.1.9) was adopted to construct maximum-likelihood trees, and based on the tree topologies, whether there were gene losses was confirmed.

2) Inspection in the Illumina assembly

To minimize the possibility of sequencing and assembly errors during PacBio data analysis, which led to false positives of gene loss, we also inspected the gene loss data in the Illumina assembly.

a Exonerate[16] (v2.2.0) software was used (--model protein2genome --percent 50) to map the protein sequences of *Arabidopsis* orthologs to *Cuscuta australis* Illumina assembly, and the target loci were annotated with Genewise[13] (v2.2.0) to obtain protein sequences.

b Each *Arabidopsis* protein sequence was set as a query to perform blastp[8] (v2.4.0) against the gene models of all the seven autotrophic species (E-value=1e-5). The subject sequences and the Genewise-annotated *Cuscuta australis* protein sequences were analyzed through the pipeline described in 1b) of this Section.

**Supplementary Note 4.**

**Whole genome duplication analysis**

*Coffea canephora* is a plant that never underwent WGD after the ancient gamma event. To gain insight into the evolution of *Cuscuta australis* genome after the gamma event, the orthologous pairs of *Cuscuta australis*, *Ipomoea nil*, and *Coffea canephora* genes in the identified orthogroups (these gene pairs were derived from speciation and/or gene duplication after the gamma event) were fed to MCScanX_h[15] (version 201610) for construction of the post-gamma-event syntenic blocks (Supplementary Fig. 6).

We next anchored every syntenic block to the chromosomes of *Coffea canephora*. There are 21969 *Coffea canephora* genes that were found to be covered by at least one syntenic block. Thereafter, we counted the numbers of the syntenic blocks from *Cuscuta australis* and *Ipomoea nil* covering each individual *Coffea canephora* gene from these 21969 gene loci, so that we could obtain the syntenic coverage information, which is described by "number of syntenic block of *Cuscuta australis*:number of syntenic block of *Ipomoea nil*:1" for each *Coffea canephora* gene locus. Among 21969 *Coffea canephora* gene loci, 4943 (22.4%) and 4604 (21.0%) showed syntenic coverage of 3:3:1 and 2:3:1, respectively (Supplementary Fig. 3 and Supplementary Table 7), and these two types were the most common ones. These data suggest that there was a genome triplication event in both *Cuscuta* and *Ipomoea* genome after the gamma event. These two most common types of syntenic blocks were anchored to their corresponding *Coffea canephora* gene loci, and positions of these *Coffea canephora* loci were plotted over the chromosomes (Supplementary Fig. 4), and they appear to scatter relatively evenly across the whole *Coffea canephora* genome, suggesting that the genome triplication event occurred at the whole genome level rather than on certain chromosomes or specific chromosome regions (Supplementary Fig. 4). A synteny circos figure was drawn using the Circos[19] (v 0.69) tool for the biggest syntenic blocks between *Cuscuta australis*, *Ipomoea nil*, and *Coffea canephora* chromosome 2 (Fig. 1c).

To investigate whether the triplication event is shared by *Ipomoea* and *Cuscuta*, we chose 574 *Coffea canephora* genes that have at least two ortholog genes in both *Ipomoea nil* and *Cuscuta australis* among all the homologous genes in the syntenic blocks. For these 574 *Coffea canephora* genes, each gene was grouped with its orthologs, and these groups were used to construct phylogenetic trees. For these trees, we assumed two models (Supplementary Fig. 5): Model A: WGD happened before the speciation between *Ipomoea* and *Cuscuta*; Model B: WGD happed independently after the speciation between *Ipomoea* and *Cuscuta*. If model A outperforms, orthologous gene pairs from *Cuscuta australis* and *Ipomoea nil*, respectively, would be grouped together; otherwise, paralogous genes of *Cuscuta australis* and *Ipomoea nil* would be grouped separately. We tested those two models by searching the 574 phylogenetic trees for clades that fit into different models (bootstrap > 0.7). Only 36 clades supported Model B, while Model A was supported by 1003 clades.

Considering the results from phylogenetic and syntenic analyses, we concluded that a common triplication event took place before the speciation between *Cuscuta* and *Ipomoea*.

**Supplementary Note 5.**

**Transcriptome analysis**

1. Transcriptome assembly and expression level analysis

Reads from *Cuscuta australis* transcriptomes and *Cuscuta pentagona* transcriptome [2] were first aligned against the corresponding host genome sequences using bowtie2[20] (v2.2.4) to remove possible host mRNA contamination. The filtered reads were aligned against *Cuscuta australis* genome independently using HISAT2[21] (v2.0.5) with parameters "-mp 3,1", achieving an average ratio of the mapped reads of 96% and 75% (Supplementary Table 8). Unmapped RNA-Seq reads of *Cuscuta australis* were extracted and de novo assembled using Trinity[22] (r20140717). These assembled transcripts were used to support gene loss analysis (see Supplementary Note 4). Transcript abundances were calculated using StringTie[23] (v1.3.0). Genes, whose average FPKM values at least 1-fold greater in prehaustoria/haustoria than in other organs, were considered to be principally expressed in prehaustoria/haustoria (Supplementary Data 3f).

The transcriptome datasets were retrieved from the Tomato Functional Genomics Database (http://ted.bti.cornell.edu/cgi-bin/TFGD/digital/sample.cgi?ID=D004), including data from tomato roots, fruits, leaves, seeds, flowers, and buds. *Ipomoea nil* transcriptome datasets (embryos, flowers, leaves, roots, seeds, and stems) were obtained from DDBJ (ftp://ftp.ddbj.nig.ac.jp/ddbj_database/dra/fastq/DRA002/DRA002647) in raw data format, and these data were processed with HISAT2 and StringTie as describe above to acquire transcript level values.

2. Identification of principally expressed tissues for all orthogroups in *Cucuta pentagona*, *Solanum lycopersicum*, and *Ipomoea nil*

A method based on Yang et al.[24] was developed to identify principally expressed tissues (PETs). Using the orthogroup information (Supplementary Data 1b) and tissue-specific transcriptomes of these three species, firstly we searched the transcriptome data for identification of the principally expressing tissues of all the genes in these orthogroups. For each orthogroup, the transcript levels of these three species' orthologs in their respective different tissues of *Cucuta pentagona*, *Solanum lycopersicum*, and *Ipomoea nil* were assigned according to their respective transcriptome data; in case of more than one gene member exist in a species in the orthogroup, the highest transcript levels were used, and the transcript levels in different tissues within a species were compared; in a given tissue, if the orthologue's transcript level is at least 1-fold greater than the average levels in all the other tissues, this tissue was assigned as the principally expressing tissue for this specific orthologue (orthologues, in case of more than one orthologue of a species exist). This was done for all the orthogroups.

Then we searched all the orthogroups for the *Cuscuta* orthologs whose principally expressing tissue is prehautoria/haustoria (1299 orthogroups were found), and the corresponding *Solanum lycopersicum* and *Ipomoea nil* orthologs' principally expressing tissues were recorded. Finally, the principally expressing tissue of these 1299 orthogroups in *Solanum lycopersicum* (505 orthogroups) and *Ipomoea nil* (515 orthogroups) was found to be root (Supplementary Data 1d).

The orthogroups whose *Cuscuta* members are missing were used to analyze the principally expressed tissues of the *Solanum lycopersicum* and *Ipomoea nil* orthologs, in order to gain insight into the expression patterns of the corresponding orthologs of the lost genes in these autotrophic plants. Finally, the principally expressing tissues of 1402 orthogroups, which do not have *Cuscuta* members (gene loss in *Cuscuta*), in *Solanum lycopersicum* and *Ipomoea nil* are listed in Supplementary Data 1d.

**Supplementary Note 6.**

**Analysis of *R* genes, *TPS*s, *P450*s, and *RLK*s.**

*R* (*resistance*) genes are critical for plant immunity. To discover *R* genes in *Cuscuta australis* genome, we first screened for the presence of NB-ARC domain (PF00931) with HMMER[25] (version 3.0), resulting in a total of 15 *R* genes in *Cuscuta australis*. Identification of NB-ARC domain was also done for the 7Ref-Species to identify the *R* genes in these reference species (Supplementary Data 3b).

Terpenes are important chemicals which are involved in plant defenses against insects and pathogens. Similarly, we identified a total of eight *terpene synthase* (*TPS*) genes, discovered by requiring the presence of both the N-terminal domain PF01397 and Cterminal domain PF03936 in the *Cuscuta australis* genome. We found 9 *TPS* genes in *Cuscuta australis*, and the same method was also applied to search for *TPS*s the 7Ref-Species (Supplementary Data 3c).

P450s are monooxygenases, which are critical for metabolism, including production of various plant secondary metabolites. P450 genes were identified using PFAM with PF00067 using HMMER[25] (version 3.0). A total of 89 *Cuscuta australis* P450 genes (Supplementary Data 3d) were identified in *Cuscuta australis* genome, and this is much less than in the 7Ref-Species (Supplementary Data 3d).

Receptor-like kinases (RLKs) are one of the largest gene families in the plant kingdom, constituting of a predicted signal sequence, single transmembrane region, and cytoplasmic kinase domain[26,27]. They play important roles in plant growth, development, and stress responses[28-30]. In *Arabidopsis thaliana*, more than 600 members of RLKs have been reported[31], part of them are functionally characterized, such as *FLS2*[32] and *CLAVATA1*[33].

A published complete list of *Arabidopsis RLK*s (610 genes) were used as the queries [31] to search in the gene families obtained from gene cluster information of *Cuscuta australis* and 7Ref-Species. We identified 339 *RLK* genes in the *Cuscuta australis* genome, which is much less than in the 7Ref-Species (Supplementary Data 3e).

**Supplementary Note 7.**

**Test of positive selection and relaxed purifying selection.**

HYPHY package[34] (v2.3.7) was selected for testing positive selection[35] and relaxed purifying selection[36].

1. Testing positive selection:

For each orthogroup, clustalW[37] (v2.1) was used to achieve multiple sequence alignment for all protein sequences, and Fasttree[18] (v2.1.9) was employed to reconstruct the phylogenetic tree. The backtrans module in the TreeBeST[1] (v1.9.2) was then used to align nucleotide sequences based on protein alignments. The phylogenetic trees and nucleotide alignments were input into aBSREL[35] to test positive selection in all branches of the phylogenetic trees. The branches which were significantly (Corrected P-value $\leq$ 0.05) positively selected were acquired. For a given gene, if the positively selection was not specific to *Cuscuta* or *Utricularia*, namely, positive selection could also be found in ancestral branches, this gene was no longer considered to be positively selected. The positively selected genes were further filtered: If any of them have values of $K < 1$ and $P < 0.05$ in the test of relaxed purifying selection (see below), they were removed from the positively selected genes. The full list of positively selected genes can be found in Supplementary Data 3g

2. Testing relaxed purifying selection

The phylogenetic trees and nucleotide alignments from above were fed to RELAX[36]. The General Descriptive model embedded in RELAX was used to estimate K values for all the branches in the phylogenetic trees. The branches with the K values smaller than 1 were considered to have potentially experienced relaxed purifying selection. Each of these branches was further treated as the foreground branch and the rest branches were set to be the backgrounds, and K and P values were recalculated. All the branches with $K < 1$ and $P < 0.05$ were considered to have experienced relaxed purifying selection. For a given gene, if the relaxed purifying selection was not specific to *Cuscuta* or *Utricularia*, namely, relaxed purifying selection could also be found in the ancestral branches, this gene was no longer considered to have experienced relaxed purifying selection. Full list of relaxed purifying selected genes can be found in Supplementary Data 3h.

**Supplementary Note 8.**

**Photosynthesis measurements**

1. Chlorophyll fluorescence measurement

The values of electron flow through PSII (ETR(II)) was measured with an imaging fluorometer (the Imaging PAM M-Series Chlorophyll Fluorescence System, Heinz-Walz Instruments, Effeltrich, Germany) connected to a computer with the control software. ETR(II) was calculated according to Genty et al. (1989) and Schreiber et al. (1995). To develop light response curves, the plants were light-adapted for at least 20 min. *C. australis* showed very low ETR(II), and the values were much lower than those of *Nicotiana tabacum*.

2. Gas exchange measurement

All measurements of gas exchange were performed on clear days by clamping one or several stems from *Cuscuta australis* into the chamber of GFS-3000 system (Portable Gas Exchange Fluorescence System GFS-3000, Heinz-Walz Instruments, Effeltrich, Germany). For gas exchange measurements, relative humidity was maintained at approximately 60%, leaf temperatures at 25 °C and $CO_2$ concentration of 400 μmol $mol^{-1}$ for *C. australis*. Light-saturated photosynthetic rates at photosynthetic photon flux density intensities form 0 to 600 μmol $m^{-2}$ $s^{-1}$ were measured. The light response curves at photosynthetic photon flux density intensities from 1000 to 0 μmol $m^{-2}$ $s^{-1}$ were also determined. However, the photosynthetic rates of *C. australis* were so low that we could not obtain stable and reliable data after several attempts (the values are far below the detection limit).
